# Uncovering Host-Parasite Dynamics: Gene Expression Shifts in *Hematodinium*-infected *Chionoecetes bairdi* in Response to Temperature Change

**DOI:** 10.1101/2025.06.05.658092

**Authors:** Aspen E. Coyle, Samuel J. White, Grace Crandall, Pam Jensen, Steven Roberts

## Abstract

Parasites can have profound effects on their hosts, and those effects can be altered by changing environmental conditions. The dinoflagellate *Hematodinium* sp. is a common and deadly parasite of the crab *Chionoecetes bairdi*, a species vulnerable to rising ocean temperatures. To examine the impact of parasitism under various temperature conditions, infected crabs (n = 9) were held under three temperature regimes (4°C, 7.5°C, and 10°C) for 17 days. RNAseq was performed on samples from three timepoints, and the relationships of temperature and time to gene expression were examined. Transcriptomes for *C. bairdi* and Alveolata symbiotes were created, and genes linked to immune function were identified within both host and parasite. Within the host, 1721 contigs were differentially expressed in response to a temperature increase, with 86% of these increased in expression. In total, 3013 contigs linked to temperature response were identified. Additionally, numerous changes in biological processes were observed in *Hematodinium* over the course of the experiment, including development and microtubule-based processes and ribosomal assembly. Through understanding the impact of changes in temperature on gene expression within both *Hematodinium* and infected *C. bairdi*, we provide a more complete picture of the response of these species to rising ocean temperatures.

## Introduction

Parasites impact their hosts in a wide variety of ways, and can thus play a number of ecological roles. In addition to altering the behavior and physiology of the host, parasites can shift competitive balance within the host. This can make the host more susceptible to additional infections, but can also reduce susceptibility. Humans infected by parasitic helminths, for example, are less prone to infection by *Giardia* (Martin et al. 2013). Parasites can also shift community composition, as has been observed in Britain, where native red squirrels are declining partly due to a virus carried by invasive and immune grey squirrels (Tompkins et al. 2003). Finally, parasites can change the composition of ecosystems. After the eradication of rinderpest in Africa the population of wildebeest exploded. The increased grazing sharply reduced fires, which resulted in increased tree cover (Holdo et al. 2009). Clearly, understanding parasites - particularly highly pathogenic ones with common hosts - can be crucial to understanding the dynamics of an ecosystem (Wood & Johnson 2015).

The parasitic dinoflagellate *Hematodinium* is a host generalist, infecting over 40 species of crab, shrimp, and lobster, including many important species for commercial fisheries and aquaculture (Li et al. 2021). Outbreaks have occurred globally, often causing major economic damage (Li et al. 2021). Prior to 1985, only six studies described *Hematodinium* infections, all of which were confined to France and the east coast of the United States (Morado et al. 2011). In the following decades, *Hematodinium* was observed throughout the North Atlantic, North Pacific, China, and Australia (Small 2012). Today, new hosts and ranges are regularly reported (Li et al. 2021; Ryazanova et al. 2021). In some host-parasite systems its prevalence is correlated with a warming climate, while in others no such correlation appears (Morado et al. 2011). Within many hosts, *Hematodinium* prevalence varies seasonally (Eaton et al. 1991; Messick 1994; Hamilton et al. 2009; Davies et al. 2019).

*Hematodinium* has an exceptionally complex life cycle, with in vitro experiments identifying at least 10 distinct stages (Li et al. 2011). Heavily infected host individuals often produce large numbers of dinospores (Li et al. 2010), which are presumed to be the infective stage. Infections occur predominantly through waterborne transmission, though the specific method of entry into the host is unknown (Shields et al. 2017). In numerous host species, including *Chionoecetes* spp., infection is closely associated with molting (Shields et al. 2007, Messick 1994; Meyers et al. 1990), with speculation that small cracks in the integument of a freshly molted crab allow entry of *Hematodinium* dinospores (Meyers et al. 1990). Upon entering the host, the dinoflagellate proliferates within the hemolymph and organs, eventually resulting in respiratory dysfunction, extreme lethargy, and mortality (Stentiford & Shields 2005).

Distributed along the continental shelf from Oregon to the southern Bering Sea, the Tanner crab (*Chionoecetes bairdi*) has substantial economic and societal importance (Heller-Shipley et al. 2021). *C. bairdi* is often infected by an undescribed *Hematodinium* species (Jensen et al. 2010). Infection rates vary seasonally, peaking in the late summer and early fall (Love et al. 1993). Summer prevalence can be quite high, with infection rates over 50% in portions of *C. bairdi*’s range (Bednarski et al. 2011). The progress from initial infection to mortality is slow, and takes place over a minimum of several months (Love et al. 1993). Heavy infections of *Hematodinium* sp. are marked by milky white hemolymph, an opaque white or pink coloration, and bitter, unpalatable flesh (Meyers et al. 1990).

The long duration between infection and symptoms, elaborate life cycle of *Hematodinium,* and challenges of experimentally inducing infection have hampered efforts to investigate this complex host–parasite system. However, the need to obtain answers is growing more critical. Infection rates are climbing in *C. bairdi*, particularly in the Bering Sea portion of its range, as are infection rates in its close relative *C. opilio* (NOAA 2020, unpublished data). Furthermore, much of the range of *C. bairdi* has recently been struck by anomalous heat events (Di Lorenzo & Mantua 2016, Cheung et al. 2020). Marine heatwaves are projected to increase in frequency and intensity due to anthropogenic warming (IPCC 2019). Understanding the environmental drivers of infection by *Hematodinium* is critical to proper management of this valuable fishery and for its preservation for years to come.

Previous research has characterized gene expression patterns in *C. bairdi* infected with *Hematodinium* at various temperatures (Crandall et al. 2021). However, that work has focused on examining pooled libraries of hosts, rather than tracking the response of individual crabs under different temperature regimes, and was not able to examine gene expression of *Hematodinium*. Transcriptomics provides a powerful tool to examine response of both host and parasite to changing environmental conditions. The purpose of this study was to improve our understanding of the dynamics within this host–parasite system by investigating the response of both *Hematodinium* sp. and infected *C. bairdi* to changes in temperature, and to track the progression of disease within infected crab. Specifically, we used transcriptomics to uncover overall changes in gene expression and stress response within both host and parasite, along with an examination of immune response.

## Methods

### Experimental Design

Male *C. bairdi* were collected with pots (n = 400) from Stephens Passage in southeastern Alaska in October 2017, a location with a reliably high prevalence of *Hematodinium* infection (Bednarski et al. 2011; ADF&G, unpublished data). Crabs were transported to the Ted Stevens Marine Research Institute in Juneau, AK and held in a flow-through system at the bottom temperature of Stephens Passage at time of capture, 7.5°C, for a 9-day acclimation period. At the end of this period, crabs that did not appear to have completely recovered from capture stress were discarded. The end of the acclimation period and beginning of the experiment is henceforth noted as Day 0.

A hemolymph sample (0.2ml) was drawn from the 179 remaining crabs selected and preserved in RNAlater (1200 µl). Crabs were divided into three groups, with 60 crabs in each experimental group and 59 in the control temperature treatment group. The control temperature group continued to be held at 7.5°C, while water temperature within the elevated-temperature and decreased-temperature treatment group was gradually changed over a two-day period to 10°C and 4°C, respectively. A second hemolymph sample was drawn from the 177 surviving crabs and preserved in RNAlater. Temperatures were maintained for an additional 15 days, for a total experimental duration of 17 days (Figure 1). The remaining crabs then had additional hemolymph samples withdrawn and preserved in RNAlater.

**Figure 1.**
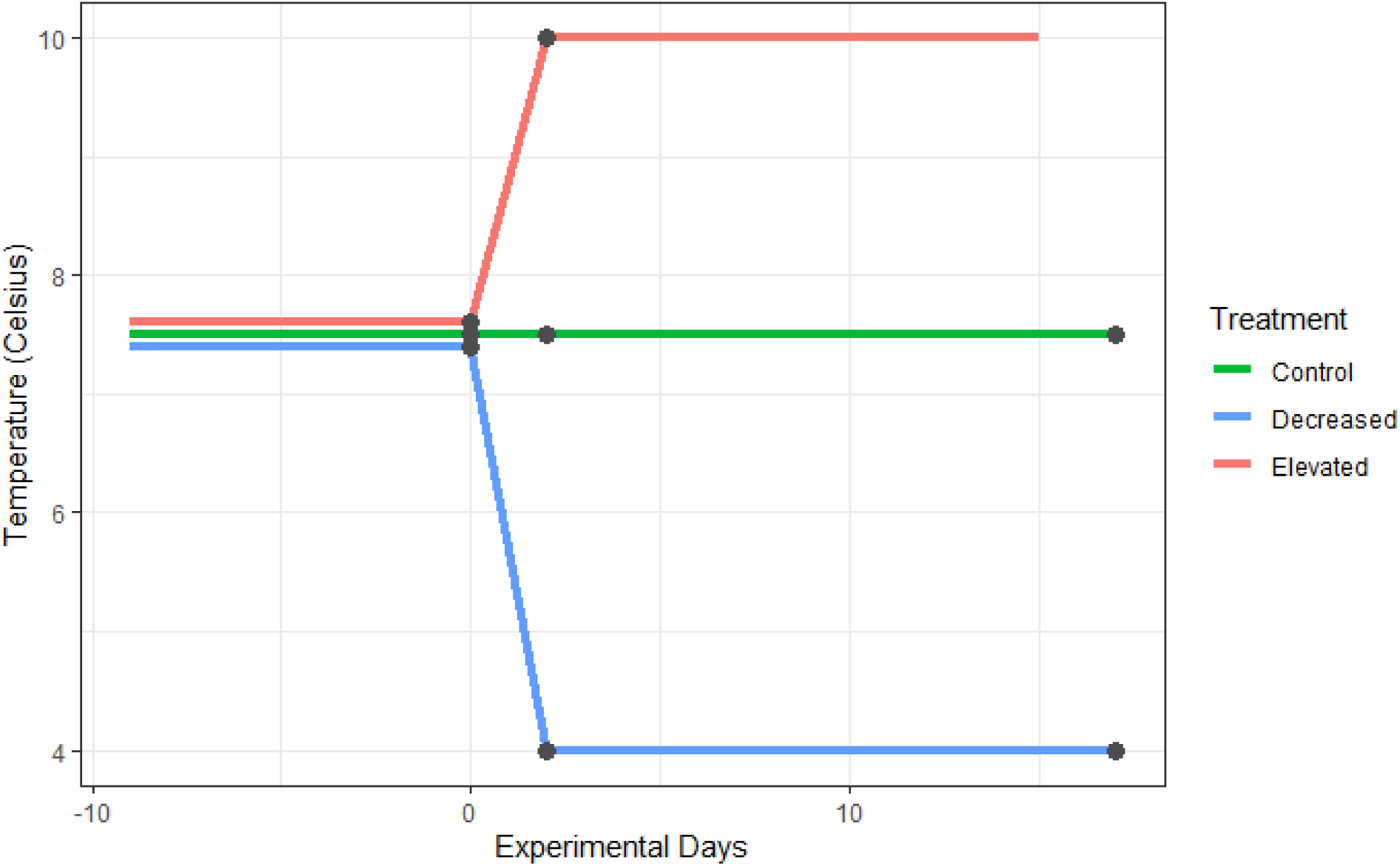
Diagram of temperature of each treatment group over the course of the experiment. Days are indexed from zero, beginning at the initiation of temperature changes for experimental groups. Three RNA samples were taken from each treatment group on days 0, 2, and 17, marked with black dots, and sequenced. Due to a mortality event, no samples with sufficiently high RNA yields were taken from elevated-temperature crabs on day 17.

The elevated-temperature treatment group saw a mass mortality event, with 58 of the 60 of crabs dying prior to the end of the experiment. Over the same period, there were eight mortalities within the decreased-temperature treatment group, and three mortalities within the control temperature group. Hemolymph samples were taken from the two surviving crabs in the elevated-temperature treatment group, but their RNA yield was not sufficient for sequencing.

### Infection Status Assessment

Hemolymph samples from the start and (if available) end of the experiment had DNA extracted, subjected to qPCR following established protocol for *Hematodinium* sp. (Crosson 2011) to determine the level of *Hematodinium* sp. infection. Samples were tested in duplicate.

### RNAseq

A total of nine crabs, three from each temperature regime, were selected based on RNA yields. As determined by qPCR, all nine were infected with *Hematodinium*. Total RNA was extracted from all hemolymph samples of these crabs using Quick DNA/RNA Microprep Plus Kit (Zymo Research) according to the manufacturer’s protocol. This created a total of 24 samples, with samples from Days 0, 2, and 17 the control and decreased-temperature treatment crabs. Due to the mortality event, the final samples were not available for elevated-temperature treatment crabs. All samples were sent to Genewiz, Inc. for library construction and RNAseq. Samples were sequenced as paired end (100bp and 150bp) on HiSeq4000 (Illumina, Inc.) sequencers.

To increase transcriptome completeness, 11 additional sequencing samples were created by pooling 112 hemolymph samples from 87 more crabs based on treatment, sampling day, and infection status (Supplemental Table 1). These samples were sent to the Northwest Genomics Center at Foege Hall at the University of Washington for RNAseq and library construction.

Samples were sequenced as paired end (100bp and 150bp) on NovaSeq (Illumina, Inc.) sequencers.

### Transcriptome Assembly and Annotation

Raw sequence data were assessed using FastQC (v0.11.8; Andrews 2010) and MultiQC (v1.6; Ewels et al. 2016) pre- and post-trimming. Data were quality trimmed using fastp (v0.20.0) (Chen et al. 2018). Trimmed reads were used for all subsequent analyses. All raw sequencing data is available in the NCBI Sequence Read Archive (SRR11548643-SRR11548677).

A transcriptome was *de novo* assembled from all individual and pooled libraries using Trinity (v2.9.0; Grabherr et al. 2011; Haas et al. 2013). This is hereafter referred to as the complete transcriptome. The complete transcriptome was assessed with BUSCO (v3.0.2; Simão et al. 2015; Waterhouse et al. 2018) using the metazoa_odb9 database, Augustus (v3.3.2; Stanke and Waack 2003; Stanke et al. 2008) with species set as fly, and hmmer (v3.2.1; hmmer.org). The transcriptome was then annotated and GO terms were obtained using DIAMOND BLASTx against the UniProtKB/Swiss-Prot database (downloaded 2021-02-09).

To examine host expression, a crab-specific transcriptome was created. To identify crab-specific sequencing reads, sequencing reads from all individual and pooled libraries were compared to the publicly available proteome (NCBI Acc: GCA_016584305.1) of a congener, *Chionoecetes opilio* (snow crab) using DIAMOND BLASTx (v0.9.29; Buchfink et al. 2015). Reads identified as matching (e-value <= 1E-04) *C. opilio* were extracted from the FastQs using seqkit (v.0.15.0; Shen et al. 2016). These crab specific reads were *de novo* assembled using Trinity (v2.12.0; Grabherr et al. 2011; Haas et al. 2013). This assembly is hereafter referred to as the *C. bairdi* transcriptome. The *C. bairdi* transcriptome was assessed for completeness with BUSCO (v3.0.2; Simão et al. 2015; Waterhouse et al. 2018) using the metazoa_odb9 database, Augustus (v3.3.2; Stanke and Waack 2003; Stanke et al. 2008) with species set as fly, and hmmer (v3.2.1; hmmer.org). The transcriptome was then annotated and GO terms were obtained using DIAMOND BLASTx against the UniProtKB/Swiss-Prot database (downloaded 2021-02-09). In another publication originating from these data, pooled libraries were aligned to the crab-specific transcriptome and analyzed for differential expression between treatment groups, as were all libraries from one of the nine crabs in this study (Crandall et al. 2022). However, analysis in that publication did not continue further to examine individual host response or non-host response.

A third transcriptome was created to examine expression in *Hematodinium* sp. Sequences from all individual and pooled libraries were taxonomically categorized with a combination of DIAMOND BLASTx (0.9.26; Buchfink et al. 2015) and MEGAN6 (6.18.3; (Huson et al. 2016)). DIAMOND BLASTx was run against NCBI nr database (downloaded 2019-09-25). The resulting DAA files were converted to RMA6 files for importing into MEGAN6 with the daa2rma utility, using the following MEGAN6 mapping files: prot_acc2tax-Jul2019X1.abin, acc2interpro-Jul2019X.abin, acc2eggnog-Jul2019X.abin. All sequencing reads categorized within and below the phylum Alveolata were identified using MEGAN6 (v6.18.3; Huson et al. 2016). Subsequently, these reads were extracted from the FastQ files using seqtk (Shen et al. 2016) and *de novo* assembled using Trinity (v2.12.0; Grabherr et al. 2011; Haas et al. 2013). Since all crabs were confirmed to be infected with *Hematodinium*, and no other Alveolata parasites of *C. bairdi* have been identified, this transcriptome likely contains only *Hematodinium* sequences. However, as the presence of other Alveolata species could not be ruled out, this is hereafter referred to as the Alveolata transcriptome. The Alveolata transcriptome was assessed for completeness with BUSCO (v3.0.2; Simão et al. 2015; Waterhouse et al. 2018) using the metazoa_odb9 database, Augustus (v3.3.2; Stanke and Waack 2003; Stanke et al. 2008) with species set as fly, and hmmer (v3.2.1; hmmer.org). The transcriptome was then annotated and GO terms were obtained using DIAMOND BLASTx against the UniProtKB/Swiss-Prot database (downloaded 2021-02-09).

This work was facilitated through the use of advanced computational, storage, and networking infrastructure provided by the Hyak supercomputer system at the University of Washington. Links to the transcriptome assemblies and files are available in Supplemental Table 2.

### Differential Expression Analysis

Quality trimmed libraries of individual crabs were pseudo-aligned to each of the three transcriptomes (complete, *C. bairdi*, and Alveolata) using kallisto (Bray et al. 2016). Two different approaches were then used to examine differential expression.

To evaluate the impact of changes in temperature on expression, the R package DESeq2 (Love et al. 2014) was used to perform pairwise comparisons. Pairwise comparisons were performed within each temperature regime, comparing expression prior to, and following, the initiation of temperature changes. Abundance matrices were produced using the Trinity (v2.12.0; Grabherr et al. 2011; Haas et al. 2013). Differentially expressed contigs, along with their accompanying accession IDs, were obtained for each comparison (Table 1).

**Table 1.** Differential expression comparisons made and the number of differentially-expressed contigs to each transcriptome.

| Temp. Regime | Comparison | Variable | DE Contigs (Complete) | DE Contigs (*C. bairdi*) | DE Contigs (*Alveolata*) |
| --- | --- | --- | --- | --- | --- |
| Elevated | Day 0 vs. Day 2 | Temp. | 367 | 1721 | 4 |
| Decreased | Day 0 vs. Day 2 | Temp. | 2033 | 7 | 0 |
| Decreased | Day 0 vs. Day 17 | Temp. | 213 | 4 | 0 |
| Decreased | Day 0 vs. Day 2+17 | Temp. | 389 | 14 | 0 |
| Control | Day 0 vs. Day 2 | Time | 7103 | 78 | 0 |
| Control | Day 0 vs. Day 17 | Time | 4764 | 473 | 7 |
| Decreased and Elevated | Day 0 vs. Day 2 | Temp. | 1113 | 192 | 0 |

In addition to pairwise comparisons within temperature regimes, a clustering approach was used to enable comparisons between treatment groups and examine correlation in expression to each variable. The R package WGCNA was used (Langfelder & Horvath 2008), which clusters contigs into eigengenes based on expression pattern and then calculates correlation between eigengene modules and experimental variables. Categorical variables were binarized, and a signed network was used. This analysis was performed once with all samples, and then again with only samples from crabs that did not die prior to the end of the experiment. This latter analysis included an examination of change in *Hematodinium* infection level over the 17-day experimental period.

### Functional enrichment

Gene ontology (GO) terms were obtained by cross-referencing the accession IDs of each contig with the Gene Ontology database. For differential expression analysis using pairwise comparisons, the log2-fold changes were extracted from the DESeq2 output and paired with GO terms as input for GO-MWU (Wright et al. 2015), which performs a Mann-Whitney U test and utilizes adaptive clustering to examine gene ontology term enrichment.

For WGCNA analyses performed with eigengene clustering, all modules with a significant correlation to a sample trait were examined, and if the significance appeared to be due to correlation to libraries from a single crab, the module was discarded. For all remaining significant modules, the module membership (kME) of its contigs was extracted, and functional enrichment of the module was analyzed using GO-MWU. This procedure was followed twice, once with all samples and once with only control and decreased-temperature treatment samples. The latter specifically examined change in *Hematodinium* infection level over time, as only one time point for *Hematodinium* infection level was available for the elevated-temperature treatment group.

## Results

### Mortality and *Hematodinium* Detection

Analysis with qPCR revealed *Hematodinium* infections were present in all crabs. Quantities of *Hematodinium* DNA were compared over the course of the experiment in control and decreased-temperature treatment groups. In four of the six crabs, *Hematodinium* infection intensity decreased, and in three it decreased by at least two orders of magnitude (Supplemental Table 3).

#### *C. bairdi* Transcriptome

Assembly of quality trimmed and crab-specific reads into a transcriptome produced 88,302 consensus sequences (Roberts, Coyle, & White 2022). A comparison against the UniProtKB/Swiss-Prot database resulted in 30,094 annotated contigs. Additional assembly statistics are in Supplemental Table 4.

#### Alveolata transcriptome

Assembly of quality trimmed reads within the phylum Alveolata into a transcriptome yielded 6,176 consensus sequences (Roberts, Coyle, & White 2022). Comparison against the UniProt/Swiss-Prot database produced 3,889 annotated contigs. Additional assembly statistics are in Supplemental Table 4.

### Immune Gene Characterization

#### C. bairdi

A number of genes within the *C. bairdi* transcriptome (n = 49) were associated with immune function (GO:0006955). Many were members of the cathepsin family, with cathepsins C, J, L, S, U, V, and W present. Cathepsin L was particularly broadly expressed, with seven distinct genes coding for cathepsin and procathepsin L. Procathepsin L was differentially expressed in the elevated-temperature treatment group over days 0 and 2. Multiple types of MAPKs (mitogen-activated protein kinases) were also present within the transcriptome, including two p38 MAPKs and one MAP4K. MAPKs are part of the IMD (immune deficiency) pathway, a notable component of the crustacean immune system. Several other genes associated with the IMD pathway were observed, including the transcription factor Relish and the kinase inhibitor IκK. NFIL3, a nuclear factor with a role in regulating Relish expression in similar systems, was also present.

Other immune-linked genes observed were Transcription Activator Protein-1 (TF AP-1) and Granzyme A. TF AP-1 acts as an immune system regulator within other crab species, along with a potential role as an osmoregulator (Wang et al. 2018). Little research on the role of Granzyme A in invertebrates has been performed, but in vertebrates it has a cytotoxic role against intracellular pathogens.

#### Hematodinium

Within the Alveolata transcriptome, four genes were linked to immune function. All four of these were cysteine proteases, which can function in blood cell degradation and invasion, surface proteins processing, and cell egress for intracellular parasites (Verma et al. 2016). Three of the four were cathepsins, including both procathepsin and cathepsin L.

### Differential Expression

Comparisons within the control temperature treatment group provided context for the frequency of differentially expressed (padj ≤ 0.05) contigs (DE Contigs) expected without a temperature change. Simultaneously, they examined expression over the course of an infection. Comparisons between Day 0 and Day 2 in an experimental group examined short-term changes to a temperature shift, while comparisons between Day 0 and Day 17 provided a long-term picture. The final comparison, between both experimental groups on Day 0 and Day 2, provided genes involved in short-term temperature response, regardless of direction.

#### C. bairdi

##### Temperature

To determine the influence of acute temperature change on gene expression in *C. bairdi*, comparisons of gene expression were made within each treatment group prior to, and two days after, the initiation of temperature changes. Within the elevated-temperature treatment group, 1721 contigs were identified as differentially expressed (padj ≤ 0.05) (Table 1). Of these, 1473 were expressed at higher levels after the increase from 7.5°C to 10°C. Within the decreased-temperature treatment group, 7 contigs were identified as differentially expressed (padj ≤ 0.05), all of which were expressed at higher levels after the decrease in temperature from 7.5°C to 4°C.

##### Time

To examine host gene expression changes as the infection develops, gene expression within the control temperature treatment group on Day 0 was compared to expression on Day 17. A total of 473 contigs were differentially expressed (padj ≤ 0.05) (Table 1). Of these, 251 were expressed on higher levels on Day 17. To determine when the changes in expression occurred, each of these groups were compared to libraries from Day 2 from the same crab. Between the first two days, there were 78 differentially expressed contigs, while the subsequent 15 days had 473 differentially expressed contigs. Functional enrichment was then examined, but no substantial enrichment in gene ontology terms was found.

#### Hematodinium *sp*

##### Temperature

To examine the impact of acute temperature change on gene expression in *Hematodinium*, the same comparisons were made with libraries aligned to the Alveolata transcriptome. Within the elevated-temperature treatment group, four contigs were identified as differentially expressed. Three of these — two (P85200 & O23717) proteasome subunits, and mitochondrial membrane ATP synthase (Q06056) — were matched to the UniProtKB/Swiss-Prot database. Over the same timeframe, no contigs were identified as differentially expressed within the decreased-temperature treatment group.

##### Time

An examination of change in *Hematodinium* expression as the infection develops was also performed by comparing expression in the control temperature treatment group between Day 0 and Day 17. A total of 7 contigs were identified as differentially expressed, and all were expressed at higher levels by the end of the experiment. When matched to the UniProtKB/Swiss-Prot database, the protein coding gene C16orf89 (Q6UX73) and a serine protease (P52717) were identified.

Significant changes in functional enrichment were observed between Day 0 and Day 17 (Figure 2). Expression decreased with time in several RNA-related processes, along with ribosomal assembly and cellular component assembly. Simultaneously, expression increased in microtubule-based processes, developmental processes, and movement of cell or subcellular components.

**Figure 2.**
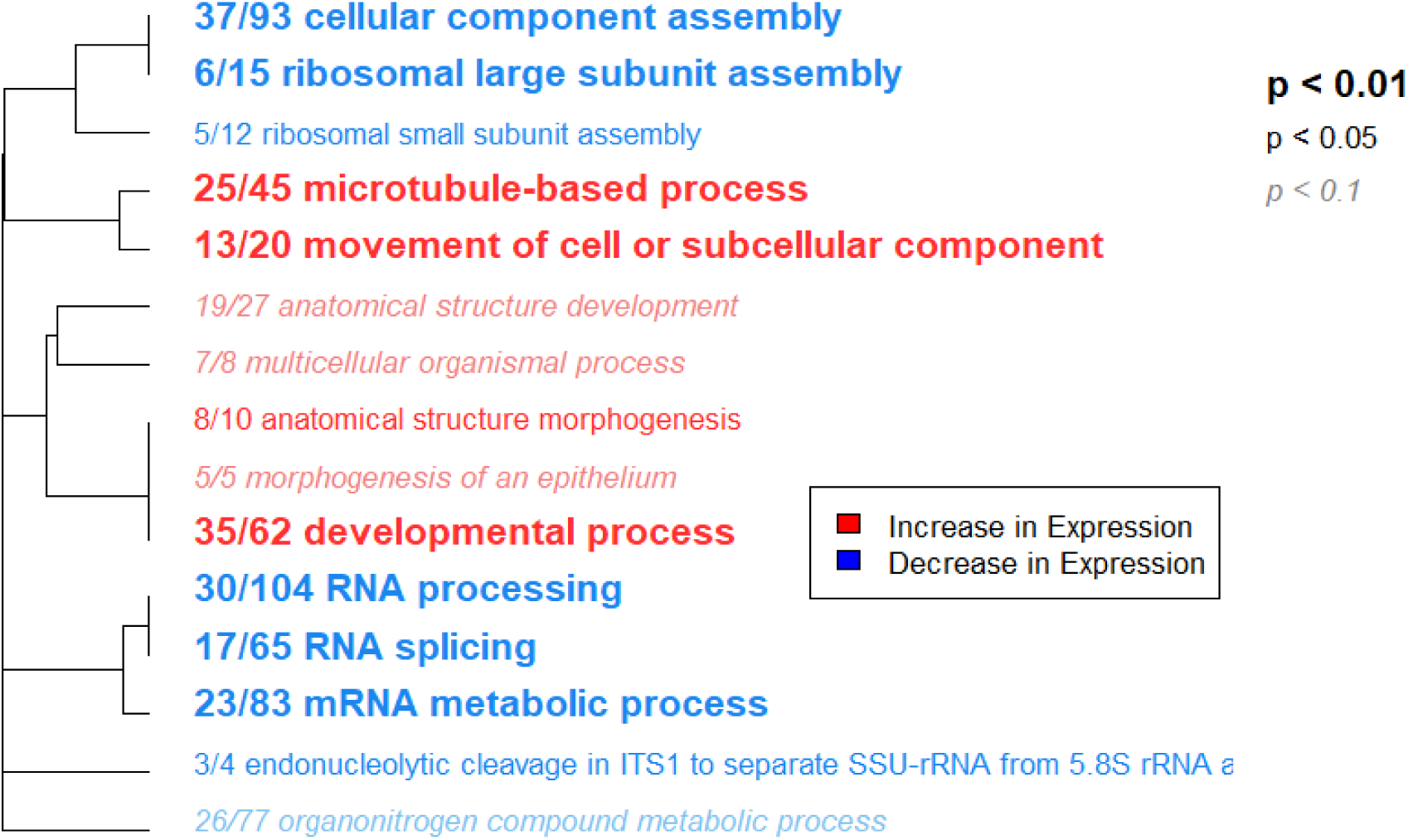
Functional enrichment of Gene Ontology (GO) Biological Process terms of parasite control temperature treatment group expression between Day 0 and Day 17 of the experiment. Tree represents hierarchical clustering based on shared genes. GO terms with zero branch length between them have gene lists in which one is a subset of the other. Text size corresponds to adjusted p-value and text color indicates the direction of regulation. Red corresponds to upregulation while blue indicates downregulation. Numbers indicate the fraction of genes with that GO term with absolute log2 fold change greater than 1.

### Clustering By Expression Patterns

#### C. bairdi

WGCNA was used to cluster genes into modules and identify their correlation with experimental variables, along with their correlation to individual crabs. Modules were named with a color to identify them. Modules with a correlation greater than ±0.5 to a single crab were not examined further, as association to a variable was likely due to expression in that individual. Two modules, black and brown, were found to be significantly linked to temperature response (Figure 3). The black module was more expressed at decreased temperatures than control temperatures (p < 0.0008), and also more expressed at elevated than non-elevated temperatures (p = 0.05). The brown module had lower expression at elevated temperatures when compared to decreased temperatures (p = 0.02), or all temperatures (p = 0.03) These modules were examined for GO enrichment, but no significant enrichment was found in either.

**Figure 3.**
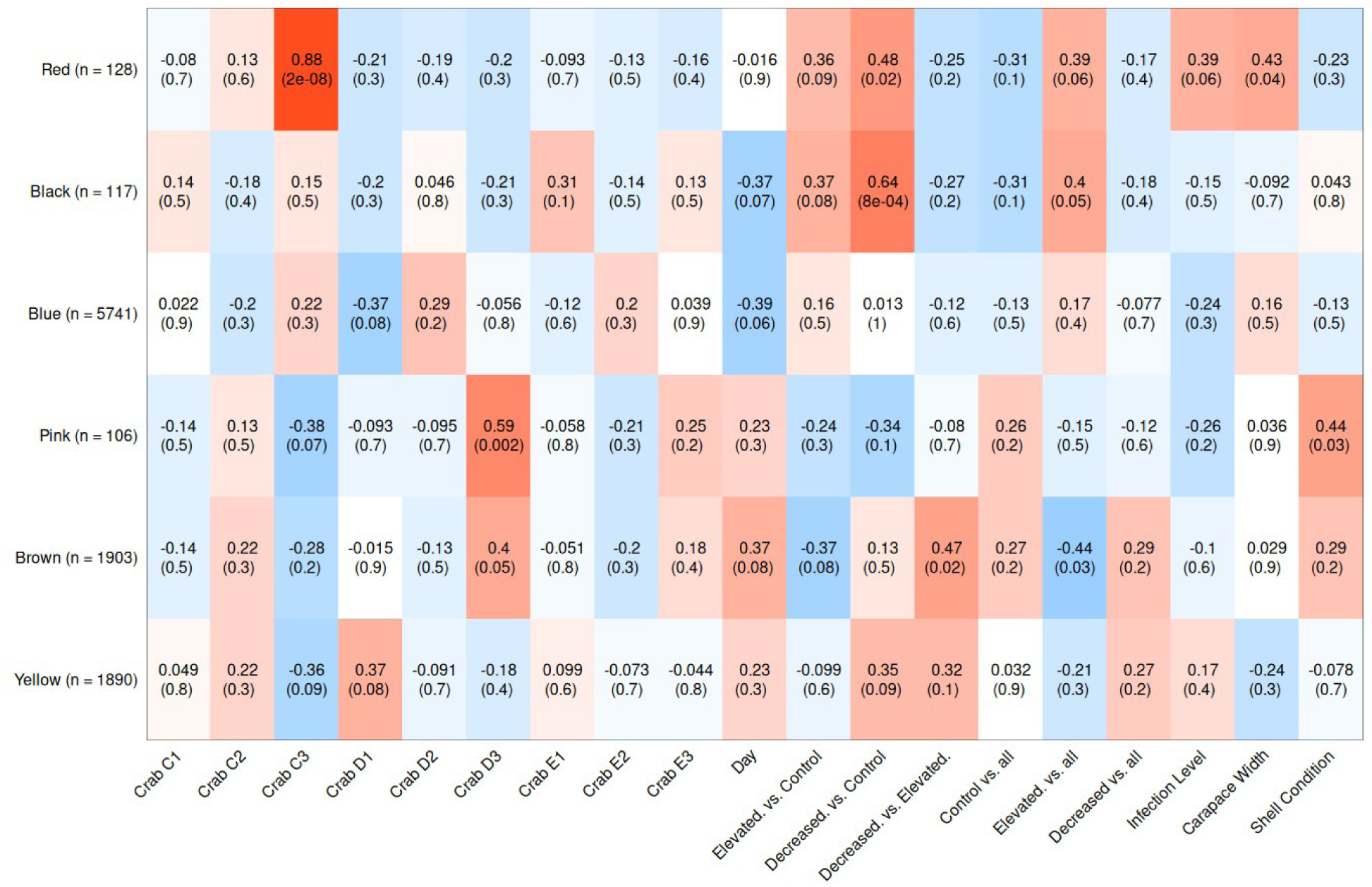
Heatmap of *C. bairdi* gene expression clusters and variables. X-axis shows variables, y-axis shows module name and the number of genes that make up the module. Each cell in the heatmap contains the correlation between the module and the variable, with the relevant p-value underneath. Cell color corresponds to correlation value, with positive correlations in red, neutral correlations in white, and negative correlations in blue.

To examine how host expression varied with change in *Hematodinium* infection over the course of the experiment, the analysis was repeated excluding samples from crabs that died prior to final sample collection. The same procedure was followed. Modules with significant correlations to variables other than change in *Hematodinium* infection were discarded, as the previous analysis was more apt for examining them. Expression in the red module (Figure 4) was significantly correlated to change in *Hematodinium* infection (p = 0.04). This module was examined for GO enrichment, but no significant enrichment was found.

**Figure 4.**
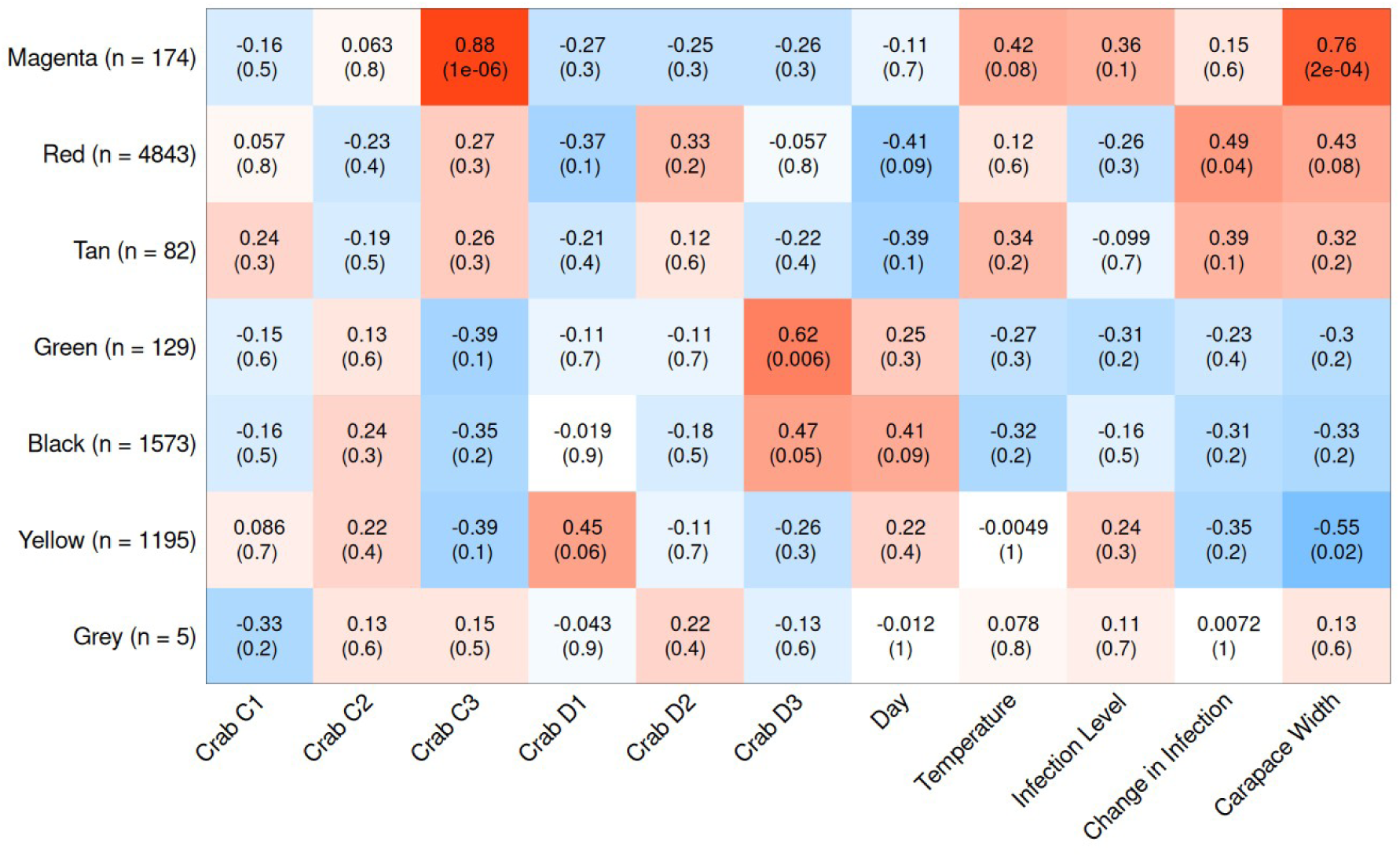
Heatmap of *C. bairdi* gene expression clusters and variables using only crabs who survived the full experiment. X-axis shows variables, y-axis shows module name and the number of genes that make up the module. Each cell in the heatmap contains the correlation between the module and the variable, with the relevant p-value underneath. Cell color corresponds to correlation value, with positive correlations in red, neutral correlations in white, and negative correlations in blue.

#### Hematodinium *sp*

The same procedure was followed for genes matching the Alveolata transcriptome. The pink module was found to decrease in expression over time (p = 0.03), while the brown module was found to be less expressed in heavily-infected crabs (p = 0.02) (Figure 5).

**Figure 5.**
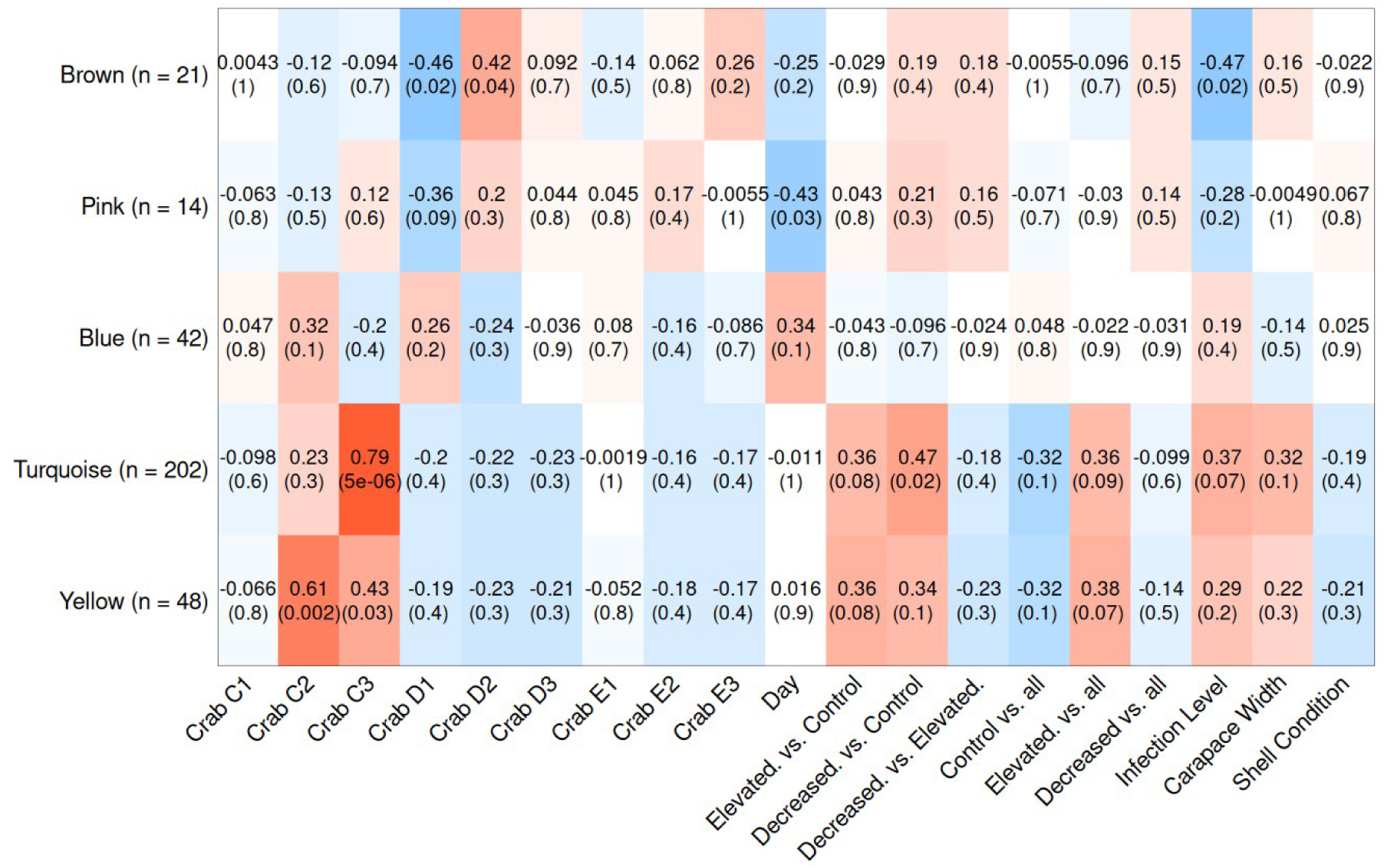
Heatmap of parasite gene expression clusters and experimental variables. X-axis shows variables, y-axis shows module name and the number of genes that make up the module. Each cell in the heatmap contains the correlation between the module and the variable, with the relevant p-value underneath. Cell color corresponds to correlation value, with positive correlations in red, neutral correlations in white, and negative correlations in blue.

The pink and brown modules were then examined for gene enrichment. Within the pink module, two pathways were enriched - negative regulation of biological processes (padj = 0.025) and cellular macromolecule catabolic processes (padj = 1 x 10^-15^). The brown module had numerous enriched pathways, including cytokinesis, vacuole organization, and translational elongation (Figure 6).

**Figure 6.**
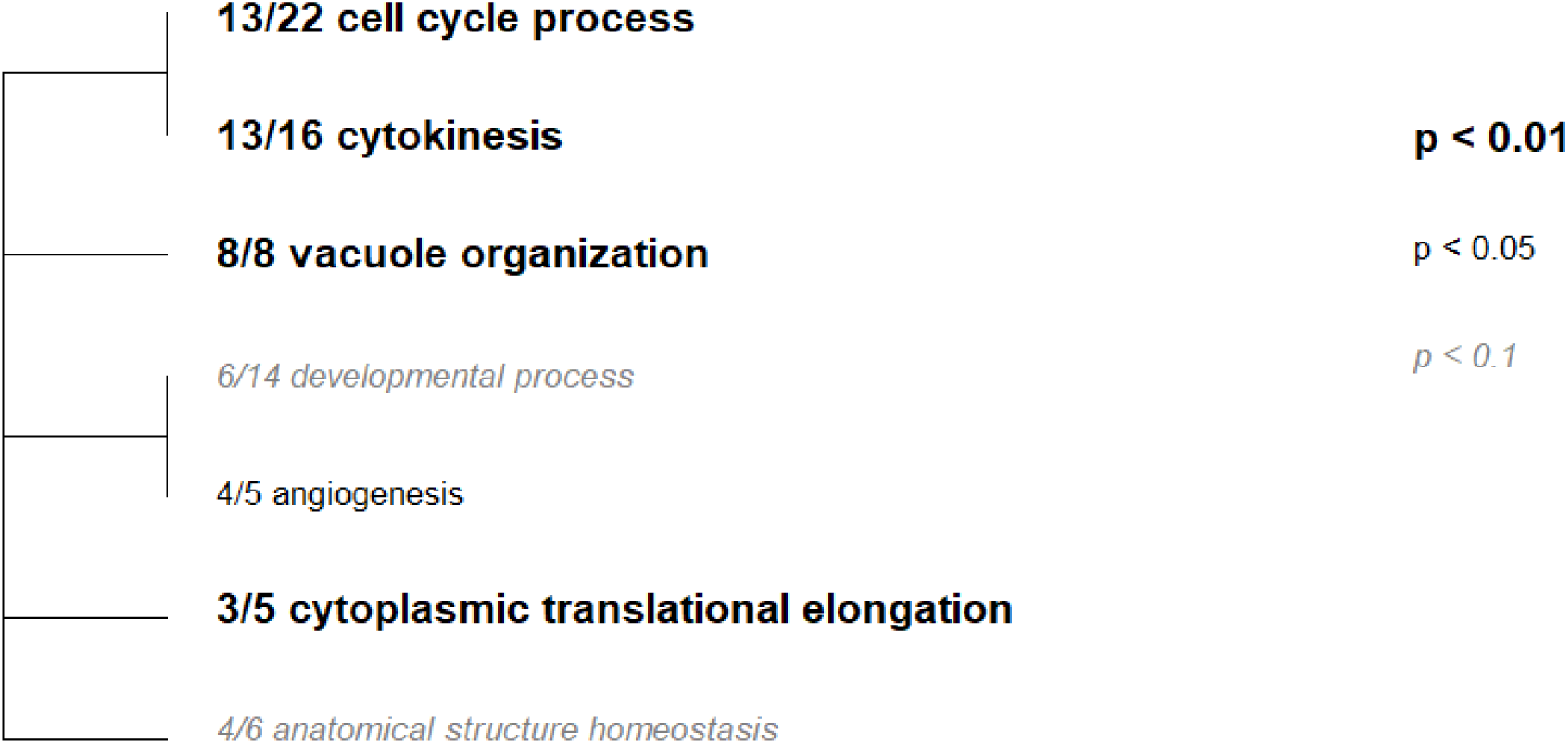
Functional enrichment of GO Biological Process terms within libraries aligned to the parasite transcriptome and clustered into the brown module. Tree represents hierarchical clustering based on shared genes. GO terms with zero branch length between them have gene lists in which one is a subset of the other. Numbers indicate the fraction of genes with that GO term with absolute log2 fold change greater than 1.

## Discussion

### Transcriptome Description

We provide a *C. bairdi* transcriptome, along with the first transcriptome for any *Hematodinium* species. We also produce the first transcriptional description of a crustacean parasitized by a dinoflagellate, providing valuable insight into the function of a host-parasite system. BUSCO scores indicate that transcriptome completeness was 73.8% for *C. bairdi* and 26.5% for *Hematodinium*.

### Immune Description

#### C. bairdi

Of the 49 immune genes observed within the *C. bairdi* transcriptome, over 20% (n = 11) were members of the cathepsin family. Seven of those code for cathepsin or procathepsin L. In multiple crustaceans, cathepsin L has been shown to be upregulated in response to pathogen exposure (Li et al. 2010) (Dai et al. 2017), indicating that further research on *C. bairdi* cathepsins may prove fruitful for uncovering the consequences of *Hematodinium* infection.

Furthermore, cathepsin C, which plays an important role in crustacean immunoregulatory function (Liu et al. 2020) was also observed within the transcriptome. Several genes associated with the IMD pathway were observed, including several MAPKs, Relish, IκK, and NFIL3.

#### Hematodinium *sp*

Four genes within the parasite transcriptome were linked to immune function, three of which were papain-family proteases. This provides an intriguing indication of the mechanism by which *Hematodinium* proliferates within the host. For species in the parasitic phylum Apicomplexa, papain-family proteases have been identified as having important roles in cell invasion (Que et al. 2002), including blood cell degradation (Pandey et al. 2005) and cell egress (Verma et al. 2016). Previous studies have found that *Hematodinium* primarily proliferates within the hemolymph of the host (Wheeler et al. 2007), but the exact mechanism of that proliferation has been undetermined. This provides an intriguing indication that papain-family proteases may play important roles in infecting and multiplying within hosts.

### Differential Expression

#### C. bairdi

Our pairwise comparison of gene expression within treatment groups identified 1721 contigs as differentially-expressed between control and elevated temperatures. Of these, 86% increased in expression following a temperature rise. This likely indicates that short-term heat exposure represents a substantial metabolic increase or stress response. Previous studies found no increase in growth rate for juvenile male *C. bairdi* raised at 6°C and 9°C (Paul & Paul 2001), suggesting that this increase in expression is largely a stress response. Since very few contigs were differentially expressed when exposed to a decrease in temperature (n = 7), this provides evidence that for infected *C. bairdi*, short-term temperature increases are much more physiologically stressful than short-term temperature decreases within the range of temperatures examined. Given the increase in frequency and severity of marine heat waves throughout much of the range of *C. bairdi* (Carvalho et al. 2021, Di Lorenzo & Mantua 2016), this indicates the substantial portion of wild *C. bairdi* populations that are infected by *Hematodinium* may experience a considerable increase in energetic costs.

The increase in temperature significantly altered expression of numerous stress-related genes. These include cytochrome p450, which is involved in detoxification in crustaceans (Steele et al. 2018), glutathione peroxidase, an enzyme that protects against oxidative stress (Cheng et al. 2020), and PAK2, which stimulates cell survival and growth (Qiu et al. 2017). Additionally, many genes with altered expression patterns were involved with both stress response and the immune system. Three chitinase genes were differentially expressed at elevated temperatures, along with two serine protease inhibitors. Studies in other crab species have shown these genes to be involved in responding to both physiological stress and bacterial infection (Zhou et al. 2018, Bao et al. 2019).

Interestingly, C-type lectin, which is a key component of the innate immune system (Zhu et al. 2016), decreased significantly in expression both after an increase in temperatures and over the 17-day course of the experiment in crabs within the control temperature treatment group. The same pattern was observed in cathepsin L, another important protein involved in the crab immune system (Li et al. 2010), and in several genes related to ubiquitin, which is involved in muscle atrophy (Koenders et al. 2002). This overlap is also seen more broadly — of the 151 identified genes that were differentially expressed between the start and end of the experiment within the control temperature treatment group, 48 were also differentially expressed within the elevated-temperature treatment group following the two-day increase in temperature. For all 48, the change in expression occurred in the same direction. The changes within the control temperature treatment group show how host expression changes as *Hematodinium* infection develops over the 17-day period. Since many of the same expression patterns were observed after only two days at elevated temperatures, this indicates that warming temperatures may speed the development of the infection. As mortality rates for *Chionoecetes* spp. infected by *Hematodinium* are remarkably high (Shields et al. 2005), heat waves could cause large mortality events.

#### Hematodinium *sp*

Within the control temperature treatment group, over the course of the experiment RNA processing and splicing decreased significantly, as did cellular component assembly and mRNA metabolic processes. Meanwhile, microtubule-based processes and developmental processes both increased in expression significantly (Figure 2).

Our clustering-based analysis of expression in all crabs also located a module of genes linked to day, with significant enrichment of cellular macromolecule catabolic processes within the module. This enrichment is possibly due to changes in dominant life stages for *Hematodinium* sp. infections. *Hematodinium* has a notoriously complex life cycle (Appleton & Vickerman 1997), with a number of different stages developing within a single host. Since we eliminated all modules predominantly linked to a single crab, this indicates that the potential change in life stage occurred within multiple hosts. Though we are unable to identify those life stages without histology samples, this provides evidence that gene expression may vary substantially between stages, and indicates that RNA sequencing may be an excellent tool for such a diagnostic.

## Conclusion

Given the economic, social, and ecological importance of *C. bairdi,* along with the prevalence and highly pathogenic nature of *Hematodinium*, research illuminating the response of this complex host—parasite system to changes in temperature is of crucial importance. Our research identified immune genes expressed by *Hematodinium* sp. along with those expressed by *C. bairdi* infected with *Hematodinium* sp., thus indicating the mechanisms through which the host defends itself and the parasite overcomes the host’s immune response. Furthermore, changes in host response under different temperature regimes, along with over the course of infection, were also identified. To better comprehend this system, future research is needed to identify the infectious stage of the parasite, determine links between expression and life stage, and more fully understand the variables associated with the development of a *Hematodinium* sp. infection. These studies would provide essential information to guide management decisions surrounding this critical resource.

## Supplemental Material

Note: Supplemental material also available at: https://github.com/afcoyle/hemat_bairdi_transcriptome/tree/main/paper/supp_files and at https://gannet.fish.washington.edu/Sebastes/

---

**Supplemental Table 1.** Description of samples pooled and sequenced to increase transcriptome completeness. Note: seven samples were present in multiple libraries.

| Treatment Group | Sampling Day | cPCR Infection | Number of Samples |
| --- | --- | --- | --- |
| Combined | 17 | Combined | 15 |
| Combined | 2 | Infected | 10 |
| Combined | 2 | Uninfected | 11 |
| Combined | 17 | Infected | 12 |
| Combined | 17 | Uninfected | 11 |
| Control | 0 | Uninfected | 10 |
| Control | 0 | Infected | 10 |
| Decreased | 2 | Uninfected | 10 |
| Decreased | 2 | Infected | 10 |
| Elevated | 2 | Uninfected | 10 |
| Elevated | 2 | Infected | 10 |

**Supplemental Table 2.** Transcriptomes created, along with links to the specific script used to create, and direct links to the file.

| Transcriptome | Lab Notation | Link to Assembly | Link to File |
| --- | --- | --- | --- |
| Complete | cbai_transcriptome_v2.0 | https://robertslab.github.io/sams-notebook/2020/05/02/Transcriptome-Assembly-C.bairdi-All-RNAseq-Data-Without-Taxonomic-Filters-with-Trinity-on-Mox.html | https://owl.fish.washington.edu/halfshell/genomic-databank/cbai_transcriptome_v2.0.fasta |
| *C. bairdi* | cbai_transcriptome_v4.0 | https://robertslab.github.io/sams-notebook/2021/03/17/Transcriptome-Assembly-C.bairdi-Transcriptome-v4.0-Using-Trinity-on-Mox.html | https://gannet.fish.washington.edu/Atumefaciens/20210317_cbai_trinity_RNAseq_transcriptome-v4.0/cbai_transcriptome_v4.0.fasta_trinity_out_dir/cbai_transcriptome_v4.0.fasta |
| Alveolata | hemat_transcriptome_v1.6 | https://robertslab.github.io/sams-notebook/2021/03/08/Transcriptome-Assembly-Hematodinium-Transcriptomes-v1.6-and-v1.7-with-Trinity-on-Mox.html | https://gannet.fish.washington.edu/Atumefaciens/20210308_hemat_trinity_v1.6_v1.7/hemat_transcriptome_v1.6.fasta_trinity_out_dir/hemat_transcriptome_v1.6.fasta |

**Supplemental Table 3.** qPCR results for each crab at the beginning and end of the experiment. NA values are due to mortality.

| Crab | Temperature Regime | Day 0 qPCR SQ | Day 17 qPCR SQ |
| --- | --- | --- | --- |
| A | Control | 283 | 1950 |
| B | Control | 316,000 | 459 |
| C | Control | 546,000 | 120,000 |
| D | Decreased | 761,000 | 6 |
| E | Decreased | 210 | 2890 |
| F | Decreased | 9140 | 32 |
| G | Elevated | 4510 | NA |
| H | Elevated | 120,000 | NA |
| I | Elevated | 211,000 | NA |

**Supplemental Table 4.** Assembly statistics for each of the three sequenced transcriptomes.

|  | **Complete** | ***C. bairdi*** | ***Alveolata*** |
| --- | --- | --- | --- |
| Number of bases of all isoforms | 819,000,346 | 44,829,042 | 4,327,777 |
| Number of isoforms | 1,412,254 | 88,302 | 6,176 |
| Number of genes | 783,006 | 47,097 | 5,395 |
| GC content | 45.41% | 52.86% | 50.22% |
| Median contig length | 325 | 317 | 582 |
| Mean contig length | 579.92 | 507.68 | 700.74 |
| N50 of isoforms (bases) | 811 | 635 | 868 |
| Number of bases of longest isoforms | 339,947,966 | 22,822,315 | 3,611,911 |
| Median contig length of longest isoforms | 285 | 294 | 548 |
| Mean contig length of longest isoforms | 434.16 | 484.58 | 669.49 |
| N50 of longest isoforms | 431 | 605 | 835 |
| BUSCO: Complete | 98.8% | 73.8% | 26.5% |
| BUSCO: Single | 24.9% | 45.8% | 20.7% |
| BUSCO: Duplicate | 73.9% | 28% | 5.8% |
| BUSCO: Fragmented | 0.9% | 7.9% | 11.2% |
| BUSCO: Missing | 0.3% | 18.3% | 62.3% |
| Annotated contigs | 147,454 | 30,635 | 53,485 |

